# Spatial Transcriptomics Analysis of Gene Expression Prediction using Exemplar Guided Graph Neural Network

**DOI:** 10.1101/2023.03.30.534914

**Authors:** Yan Yang, Md Zakir Hossain, Eric Stone, Shafin Rahman

**Author notes:** Email addresses:* (Yan Yang), (Md Zakir Hossain), (Eric Stone), (Shafin Rahman).

## Abstract

Spatial transcriptomics (ST) is essential for understanding diseases and developing novel treatments. It measures the gene expression of each fine-grained area (i.e., different windows) in the tissue slide with low throughput. This paper proposes an exemplar guided graph network dubbed EGGN to accurately and efficiently predict gene expression from each window of a tissue slide image. We apply exemplar learning to dynamically boost gene expression prediction from nearest/similar exemplars of a given tissue slide image window. Our framework has three main components connected in a sequence: i) an extractor to structure a feature space for exemplar retrievals; ii) a graph construction strategy to connect windows and exemplars as a graph; iii) a graph convolutional network backbone to process window and exemplar features, and a graph exemplar bridging block to adaptively revise the window features using its exemplars. Finally, we complete the gene expression prediction task with a simple attention-based prediction block. Experiments on standard benchmark datasets indicate the superiority of our approach when compared with past state-of-the-art methods.

**Graphical Abstract:** 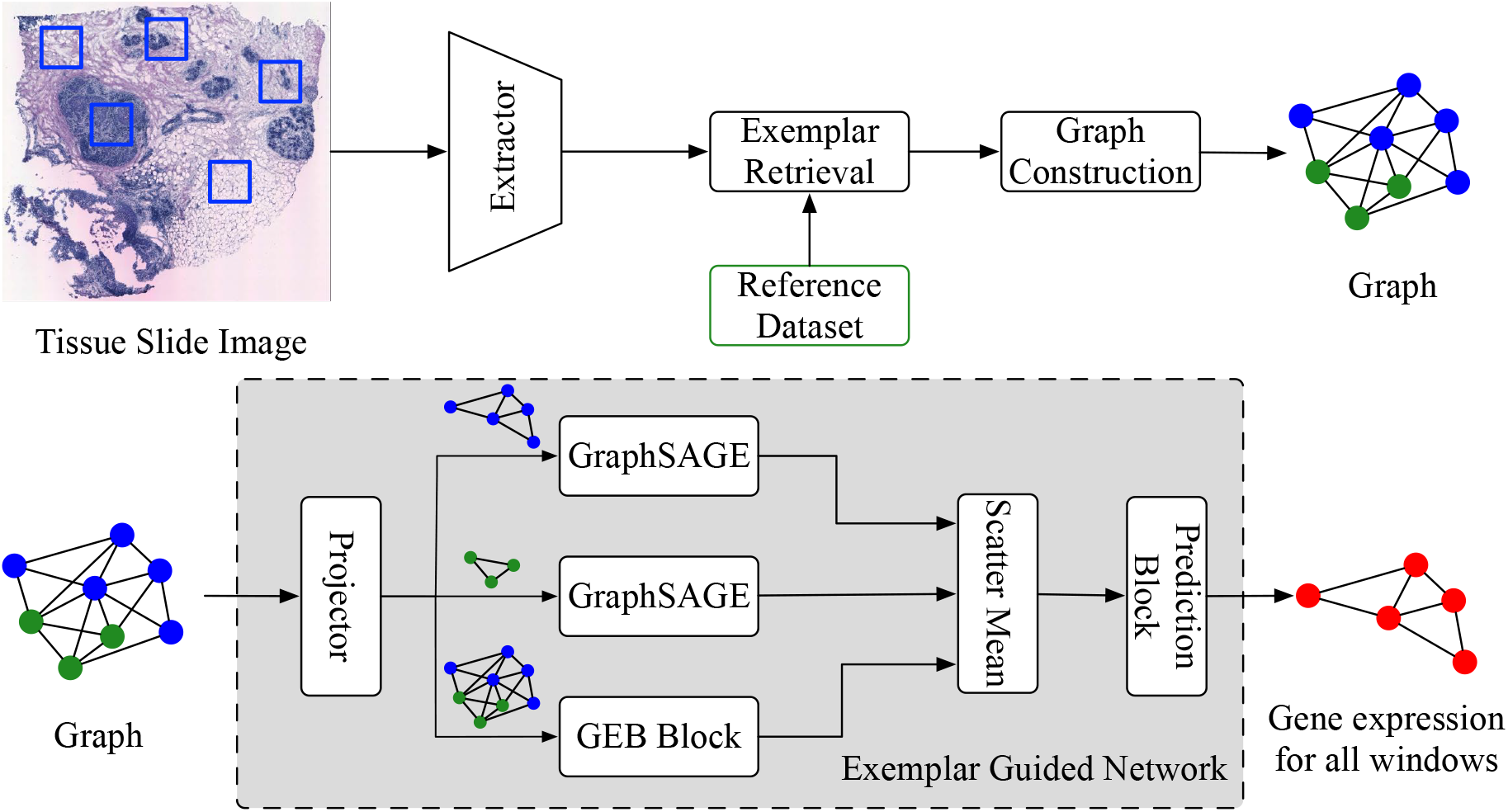

In this paper, we aim to predict gene expression of each window in a tissue slide image. Given a tissue slide image, we encode the windows to feature space, retrieve its exemplars from the reference dataset, construct a graph, and then dynamically predict gene expression of each window with our exemplar guided graph network.

**Highlights:**

- We propose an exemplar guided graph network to accurately predict gene expression from a slide image window.
- We design a graph construction strategy to connect windows and exemplars for performing exemplar learning of gene expression prediction.
- We propose a graph exemplar bridging block to revise the window feature by using its nearest exemplars.
- Experiments on two standard benchmark datasets demonstrate our superiority when compared with state-of-the-art approaches.

## 1. Introduction

Based on an editorial report of the Natural Methods [1], spatial transcriptomics (ST) is the future of studying disease because of its capabilities in measuring gene expression of fine-grained areas (i.e., different windows) of tissue slides. However, ST is in low throughput due to limitations in concurrent analysis for the candidate windows [2]. To accurately predict gene expression from each window of a tissue slide image (Fig. 1), this paper proposes a solution named exemplar guided graph network (EGGN), allowing efficient and concurrent analysis.

**Figure 1:**
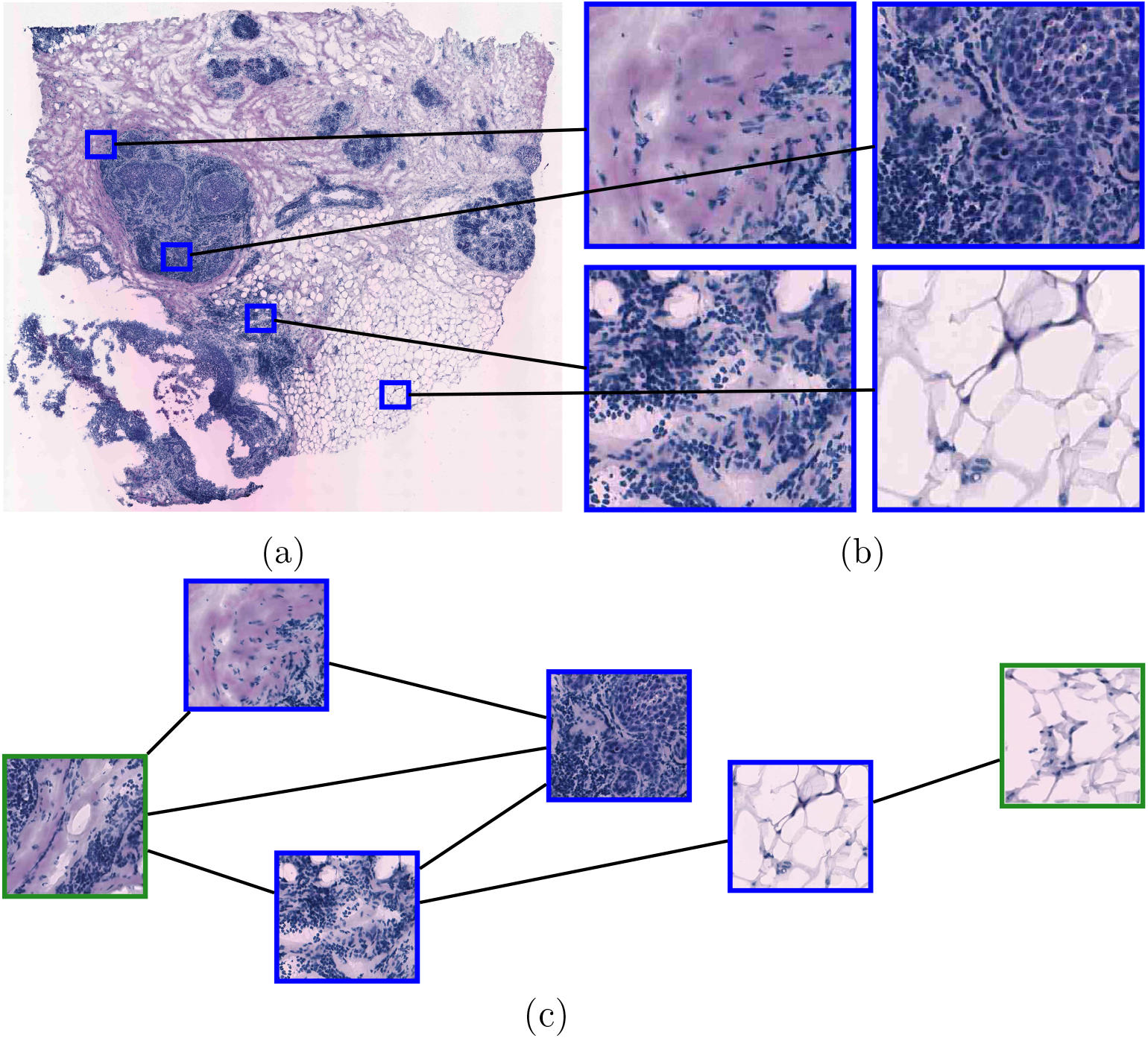
Overview of fields. Each fine-grained area (i.e., window) of (a) a tissue slide image has a distinct expression of the same gene types. For example, we have a tissue slide image with (b) four windows and each of the windows corresponds with the different expression of the same gene types. Our goal is to predict the gene expression of each window. Our key idea is to build (c) a graph connecting windows spatially close with each other and the exemplars for each window, which is used by our proposed framework to dynamically benefit gene expression prediction.

Previous works adopt end-to-end neural networks, namely STNet [3] and NSL [4], to independently establish a mapping between gene expression and the slide image window. STNet is a transfer learning-based approach that finetunes a pretrained DenseNet [5] for the gene expression prediction task. On the contrary, NSL maps the color intensity of the slide image window to gene expression by a single convolution operation. Though amenable to high throughput because of using neural networks, their prediction performance is inferior.

We investigate two important limitations of the existing approaches [3, 4]. i) Local feature aggregation: gene expression prediction can be considered as individually aggregating and identifying the feature of each gene type for the slide image window. The long-range dependency, i.e., global context, among identified features is needed to reason about complex scenarios [6, 7], as those features are generally non-uniformly distributed across the slide image (see Sec. 3 for details). STNet (i.e., a pure convolution approach) emphasizes local context during feature aggregation within a limited slide image window. It fails to bring interaction among features that are apart from each other. By experimenting with extensive state-of-the-art (SOTA) network architectures, we show that models with local feature aggregation achieve low performance when compared to models with long-range dependencies on the slide image. ii) Vulnerable assumption: identifying gene expression by directly detecting the color intensity of the slide image window is vulnerable. By experimenting on standard benchmark datasets, we show that NSL only works in extreme cases. For example, in the STNet dataset [3], tumor areas of slide image windows are usually purple, which benefits tumor-related gene expression prediction. This method finds a negative Pearson correlation coefficient (PCC) when evaluating the model with the least reliable gene expression prediction, i.e., PCC@F in Tab. 1.

Furthermore, as shown by [3], windows of a slide image spatially close with each other usually exhibit similar features, suggesting correlated gene expression among them. However, both of the above two approaches independently predict the gene expression of each window and ignore interactions among these windows. Windows distributed over the slide image can be connected to construct a graph for gene expression prediction with a graph convolutional network (GCN) [8]. Compared to independently modeling the window, graph-based modeling of the task allows reasoning dependency among windows for predicting gene expression, considering the local structure/context/spatial relations among windows.

In this paper, we propose an EGGN framework to address the above limitations. EGGN uses GraphSAGE [9] as a GCN backbone and incorporates exemplar learning concepts for the gene expression prediction task. To enable exemplar retrieval of a given slide image window, we use a feature extractor for defining a feature space to calculate the similarity between two slide image windows. Then, we construct a graph, considering the local context (i.e., nearby windows) and global context (i.e., shared exemplars act as anchors for information propagation) for the gene expression prediction task. We use GraphSAGE layers as our backbone to allow interactions among windows and exemplars. Meanwhile, we have a graph exemplar bridging (GEB) block to update window features by the exemplars and the gene expression of exemplars. Allowing dynamic information propagation, the exemplar feature also receives and is updated with the status of the window features. Semantically, the former update corresponds with ‘the known gene expression’, and the latter corresponds with ‘the gene expression the model wants to be known’. Finally, we have an attention-based prediction block to aggregate exemplars of each window and the exemplar-revised window features, for predicting gene expression.

Our contributions are summarised as follows:

- We propose an EGGN framework, a GCN-based exemplar learning approach, to accurately predict gene expression from the slide image window;
- We design a graph construction strategy to connect windows and exemplars for performing exemplar learning of gene expression prediction under GCNs;
- We propose a GEB block to revise the window feature by using its nearest exemplars;
- Experiments on two standard benchmark datasets demonstrate our superiority when compared with SOTA approaches.

A preliminary version of this paper has been published previously [11]. However, it considers interactions between exemplars and each window independently, and ignores the exemplar shared by multiple different windows potentially serve as anchors for facilitating information propagation among spatial apart windows of a slide image. In this paper, we extend our previous version as follows: i) we demonstrate that reasoning spatial relations among windows out-weights feature learning in our problem, which motivates us to design a GCN-based exemplar learning framework (Fig. 2); ii) we address the above independent interaction and information propagation bottleneck issues in our new exemplar learning framework; iii) we analyze the proposed framework with careful ablation designs, *e*.*g*., we show that the proposed exemplar learning strategy benefits general GCN frameworks; iv) we explore SOTA GCN-based methods in extensive experimental comparisons.

**Figure 2:**
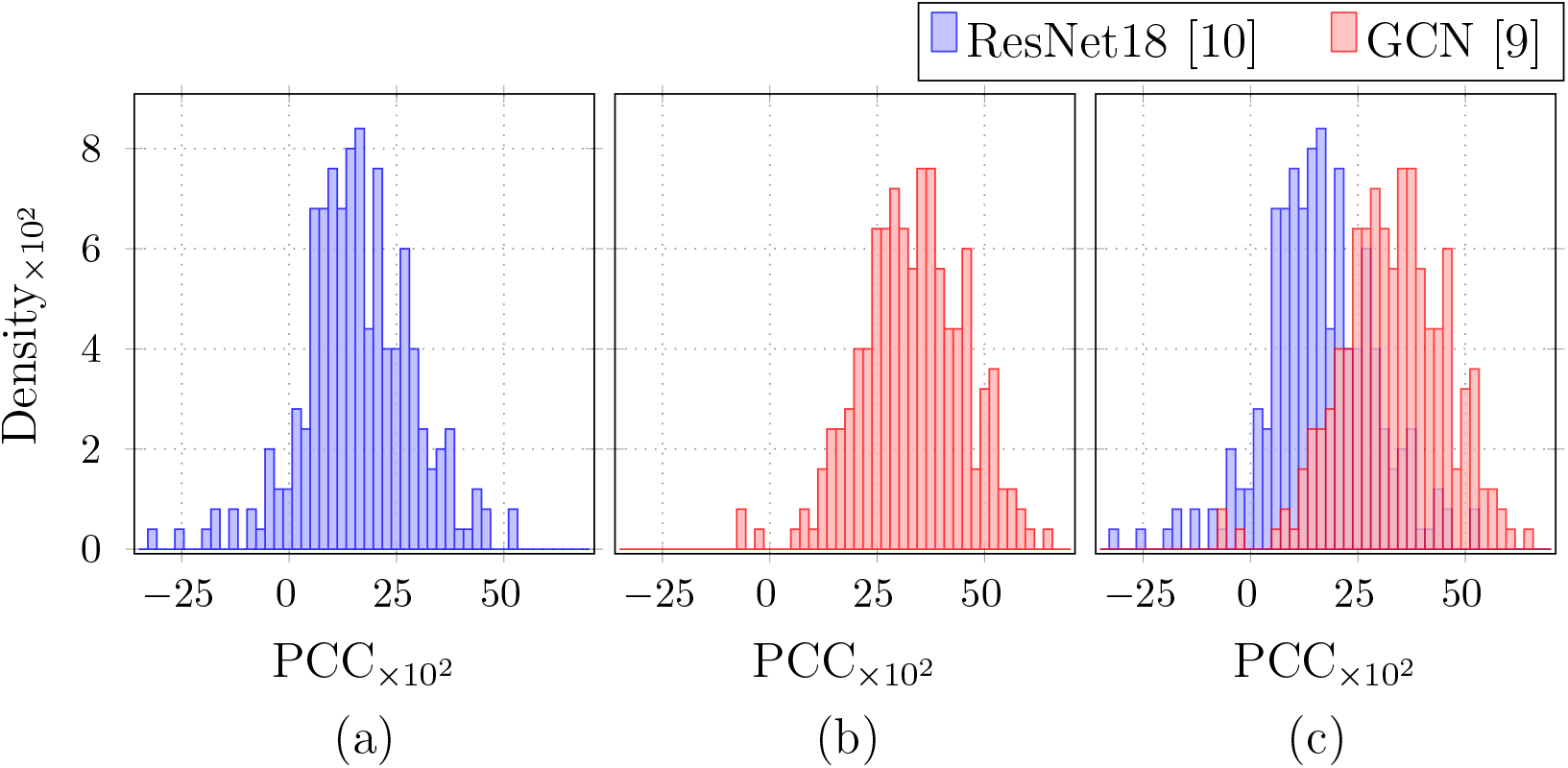
We compare the PCC distribution obtained by using (a) ResNet18 model and (b) GCN model for gene expression prediction, where window features used in the GCN [9] are extracted by the ImageNet-1K pre-trained ResNet18 [10]. The x-axis and y-axis respectively denote the PCC and the density at different PCC. We also combine the plot and plot (b) into the plot (c), for a better comparison. As shown, modeling the spatial relations among windows out-weights feature learning of each window.

## 2. Related Work

This section first reviews the study of gene expression prediction. Then, we summarise recent exemplar learning achievements in natural language processing and computer vision domains. Finally, we briefly introduce the background of GCNs.

### Gene Expression Prediction

Measuring gene expression is a fundamental process in developing novel treatments and monitoring human diseases [12]. Recently, deep learning methods have been introduced to this task. Existing methods predict the gene expression either from DNA sequences [12] or slide images [3, 4, 13]. This paper explores the latter approach which is divided into two streams. First, Schmauch *et al*.[13] employs a multi-stage method including pretrained ResNet feature extractions [10] and a K-Means algorithm to model the Bulk RNA-Seq technique [14]. It measures the gene expression across cells within a large predefined area that is up to 10^5^ × 10^5^ pixels in a corresponding slide image [13]. However, this approach is ineffective for studies that require fine-grained gene expression information, such as tumor heterogeneity [15]. Second, on the contrary, He *et al*. [13, 5] *and Dawood et al*. [4] design a STNet and an NSL framework to predict the gene expression for each window (i.e., fine-grained area) of a slide image. This corresponds with the ST technique [16]. We model ST to predict gene expression, as this potentially solves the bulk RNA-Seq prediction task simultaneously [3]. For example, aggregation of gene expression predictions for each window across a slide image results in a bulk RNA-Seq prediction.

### Exemplar Learning

The K-nearest neighborhood classifier is the most straight-forward case of exemplar learning. It classifies the input by considering the class labels of the nearest neighbors. Exemplar learning is a composition of retrieval tasks and learning tasks [17]. It has been widely employed to increase the model capability by bringing in extra knowledge of similar exemplars. The applications of exemplar learning include visual question answering [18, 19, 20, 21], language generation [22, 23, 24, 25], real-time visual object tracking [24], fact-checking [26], fact completion [27, 28], and dialogue [29]. However, most of the exemplar learning approaches do not apply to our task because of the domain shift. This paper investigates an application of exemplar learning in gene expression prediction from slide image windows. As a result, we devise a GEB block to adapt exemplar learning to our task.

### Graph Convolutional Network

GCNs are an extension of Convolutional Neural Networks (CNNs) designed for modeling non-Euclidean structured data such as graphs and reasoning the underlying complex relationships [8]. GCNs can be categorized into two main groups: spectral-based and spatial-based approaches. Spectral-based GCNs rely on the graph Laplacian and perform convolutions of spatial domain in the frequency domain [30, 31], while spatial-based GCNs utilize message-passing frameworks to progressively propagate information from source nodes to their neighbors (target nodes) along the edges of the graph. After information propagation, target nodes aggregate all received information, potentially bringing the perception of the graph structure into final node features [9, 32]. Some works also explore edge attributes during the propagation process [33, 34, 35]. GCNs with different information propagation and aggregation strategies have been proposed for diverse vision tasks [36, 37, 9, 32, 38]. In this paper, we propose a GEB to manage information propagation between windows and exemplars for the gene expression prediction task. The GEB block dynamically revises the window features using the nearest exemplars and updates the exemplars to improve the gene expression prediction.

## 3. EGGN Framework

### Problem Formulation

We have a tissue slide image containing multiple windows; each window is annotated with gene expression. We denote the slide image as pairs of slide image windows **w**_*i*_ and gene expression **y**_*i*_, i.e., 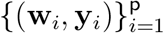, where p is the set size. We aim to train a neural network model to predict **y**_*i*_ from **w**_*i*_.

From the ST study [3], two main challenges exist. i) Long-range dependency: gene expression-related features are non-uniformly distributed over the slide image (refer to Figure 3 of [3] for evidence). The interactions among these features are needed to group expression of the same gene type; ii) Skewed gene expression distribution: the expression of some gene types has a skewed distribution, similar to the imbalance class distribution problem. This skewed distribution (see Fig. 3 for an example) poses challenges in predicting the expression of these gene types. In this paper, we attempt to mitigate them by learning from similar exemplars.

**Figure 3:**
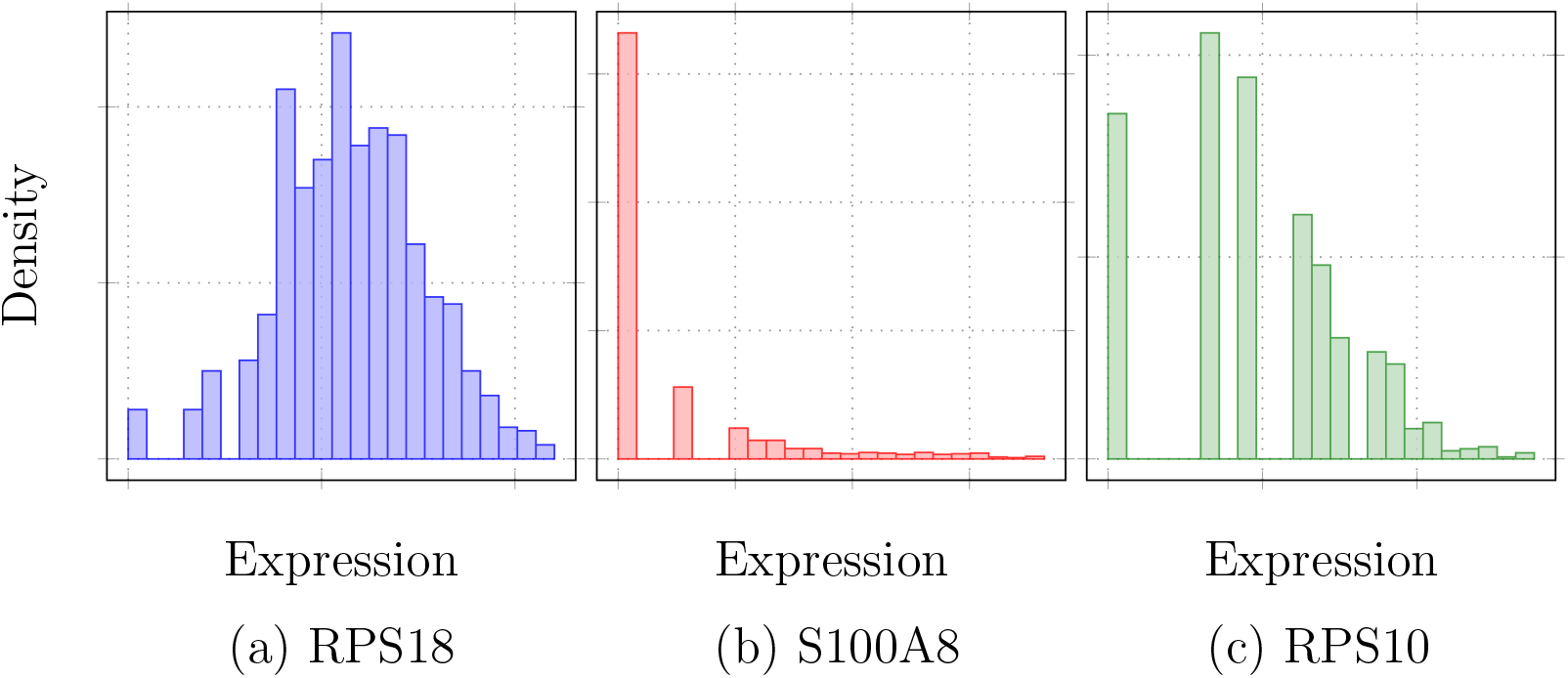
Gene expression distributions of STNet dataset [3]. Each gene expression is log-transformed. (a) is the well-distributed expression of gene RPS18. (b) and (c) are the long-tail distributed expression of gene S100A8 and gene RPS10.

### Model Overview

With these motivations, we have designed the EGGN framework containing three main modules in sequence. The overview of our frame-work is shown in Fig. 4. i) Exemplar retrieval (Sec. 3.1): we have a feature extractor **F**(·) to embed the slide image window **w**_*i*_ in to feature 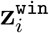, i.e., 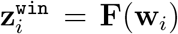. Meanwhile, there is a reference dataset 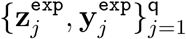 con-taining q pairs of exemplars 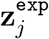 embeded by the feature extractor **F**(·) and the exemplar gene expression 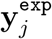. In the feature space of **F**(·), we then retrieve the nearest exemplars of **w**_*i*_ from the reference dataset to form a set 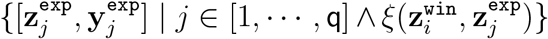, where *ξ*(·, ·) is the kNN algorithm used to determine if 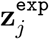 is the nearest exemplar of 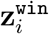. ii) Graph construction (Sec. 3.2): we construct an exemplar-based graph 𝒢 = (𝒱, ℰ) for the slide image. Each window or each exemplar can be considered as a node, and they form two sets of nodes, a window node set 𝒱^win^ and an exemplar node set 𝒱^win^. The union of 𝒱^exp^ and 𝒱 is 𝒱, i.e., 𝒱 = 𝒱^wi^ ∪ 𝒱^exp^. The edge set is denoted as ℰ = ℰ_win→win_ ∪ ℰ_exp→exp_ ∪ ℰ_win→exp_ ∪ ℰ_exp→win_, exploring relations of window to window ℰ_win→win_, exemplar to exemplar ℰ_exp→exp_, window to exemplar ℰ_win→exp_, and exemplar to window ℰ_exp→win_. In our formulation, 𝒢 is a heterogeneous graph. iii) Exemplar learning (Sec. 3.3): we train a model **C**(·, ·, ·) that maps 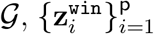, and 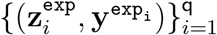 to 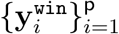 by a single foward pass. Our model uses a GraphSAGE-based backbone. We bring interactions between 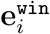 and 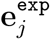 to leverage 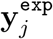 with a proposed GEB block, whenever *e*_*ji*_ ∈ ℰ^exp→win^. Meanwhile, for facilitating their interactions, the 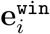 is propagated back to revise 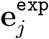. With the introduction of exemplars, our framework dynamically benefits when predicting gene expression.

**Figure 4:**
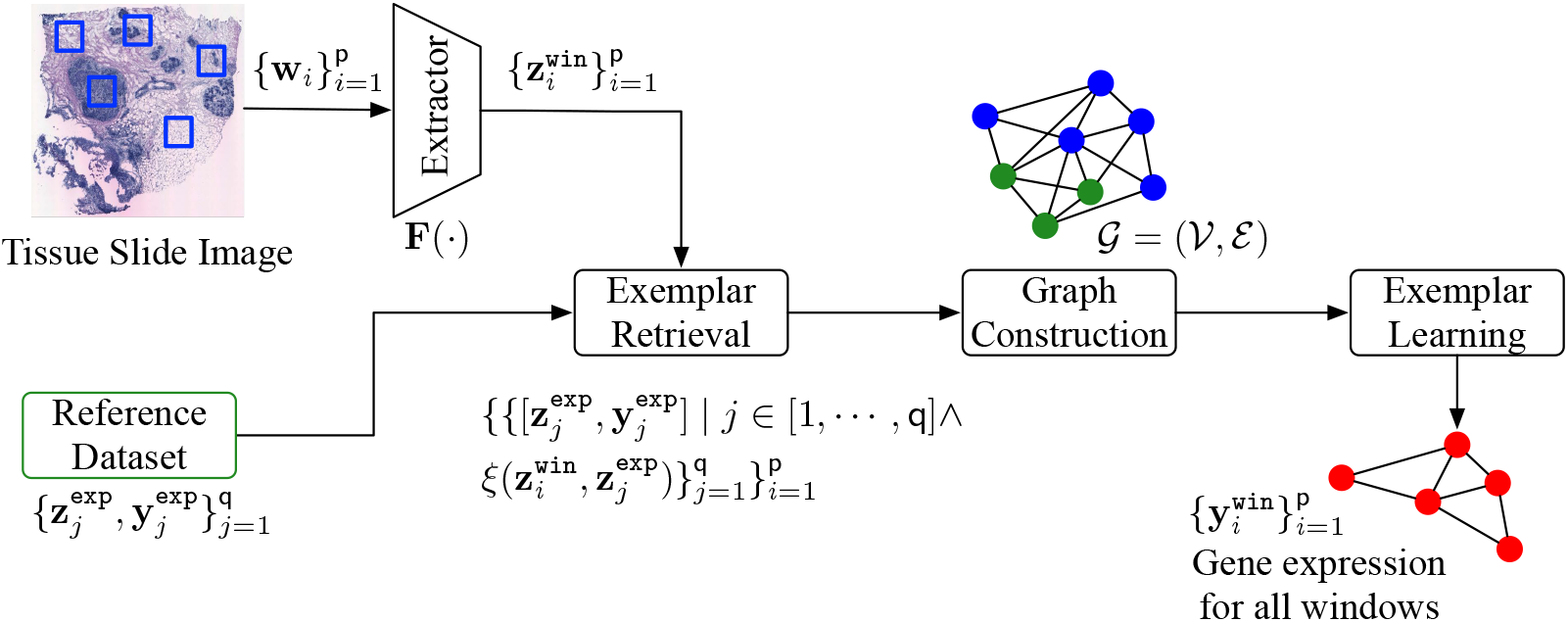
Framework overview. Given a slide images containing p windows **w**_*i*_, i.e.,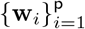, we embed each window **w**_*i*_ into features by using the feature extractor **F**(·). We have 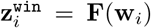. Meanwhile, there is a reference database 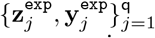 that collects q pairs of exemplar embedded by the same **F**(·) and the gene expression of the exemplar. For each window **w**_*i*_, we then perform exemplar retrieval from the reference database, resulting in 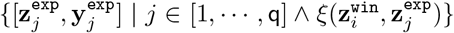, where *ξ*(·, ·) determines if 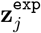 is the nearest exemplar of 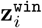. We then construct a graph 𝒢 = (𝒱, ℰ). 𝒱 is a node set of windows and exemplars, and ℰ is edges that connect the nodes. We finally perform exemplar learning on the graph 𝒢, obtaining gene expression for all window nodes, i.e., 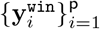.

### 3.1. Exemplar Retrieval

To retrieve the exemplar of a given window **w**_*i*_, we have an extractor (i.e., an encoder) that is coupled with a distance metric to amount the similar-ity between the given window and exemplar for a reference dataset for the exemplar retrieval.

#### Feature Extractor

Unlike our previous work [11] that trains a StyleGAN-based autoencoder [39, 40] for retrieving the exemplars, we found that an ImageNet-1K pre-trained ResNet brings a more accurate gene expression prediction in the graph-based setting. We denote **F**(·) as a ResNet feature extractor, where the classification layer is removed from a standard ResNet. Though **F**(·) is trained from a different domain, we show that **F**(·) effectively encodes textures for gene expression prediction, by comparing it with other possible encoders in a later section.

#### Method

We use the extractor **F**(·) for the exemplar retrieval. The window **w**_*i*_ is embedded into 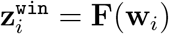. In the feature space of **F**(·), we measure the similarity between 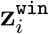 and each exemplar 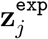 from the reference dataset by using the Manhattan distance ℒ_1_. We then execute the kNN algorithm *ξ*(·, ·) for retrieving the nearest exemplar set. We empirically verify the optimal number of used exemplars in Sec. 4.2. To generalize the model performance, we restrict that the candidate image pairs are from different patients. The overall process is presented in Fig. 5. In the experiment section, we compare the proposed exemplar retrieval with alternative retrieval approaches.

**Figure 5:**
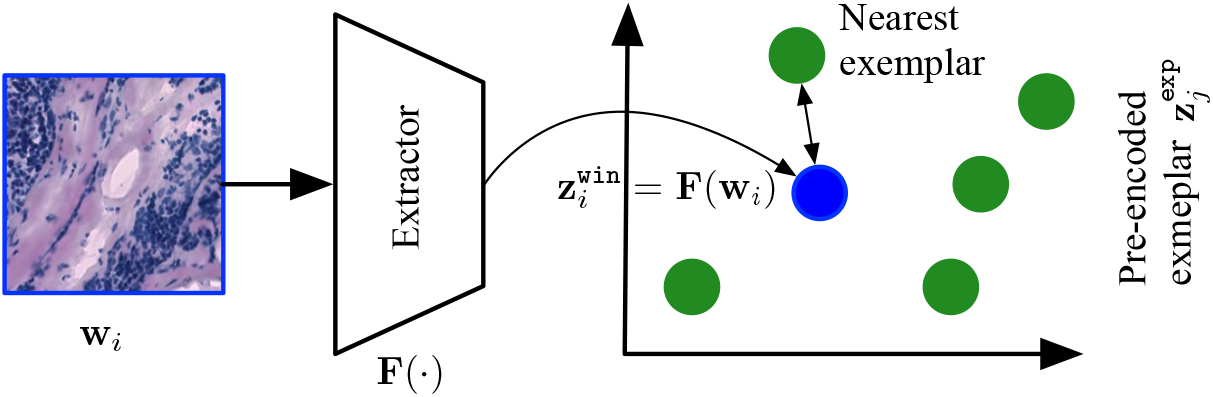
Overview of the exemplar retrieval. The green cycles are pre-embeded exemplar features, *e*.*g*.,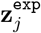. Given a window **w**_*i*_, we extract 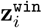 with **F**(·) and retrieve the nearest exemplar.

### 3.2. Graph Construction

We construct a graph 𝒢 = (𝒱, ℰ). Each window and exemplar is treated as a node, forming a window node set 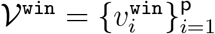 and an exemplar node set 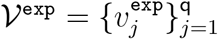. We take an union of the two set, 𝒱 = 𝒱^win^ ∪ 𝒱^exp^, to obtain a node set of our graph 𝒢. As described, we consider four edge types.

They are

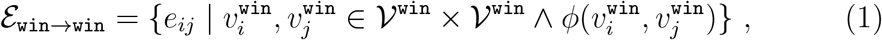

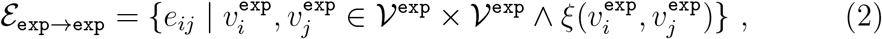

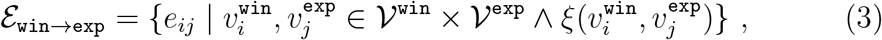

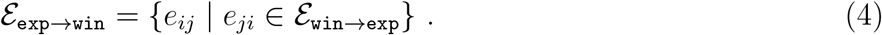

*φ*(·, ·) explores the spatial relation of windows, determining if the spatial distance of two input windows in the slide is below a given threshold. Note that *ξ*(·, ·) is a kNN-based function in the feature space of ℱ (·) for connecting k-nearest neighbors. Thus, we have the edge set of the graph 𝒢 as ℰ = ℰ _win→win_ ∪ ℰ _exp→exp_ ∪ ℰ _win→exp_ ∪ ℰ _exp→win_.

### 3.3. Exemplar Learning

Our model **C**(·, ·, ·) is composed of a projector, a *L*-layer GraphSAGE-based backbone, GEB blocks, and a prediction block, where we use the GEB block in each of the backbone layers. With graph 𝒢, our model maps node features 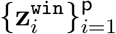 of a slide image and exemplars 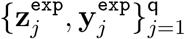 to gene expression 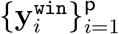 of each window. Our model allows interactions between

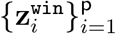 and 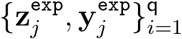 to progressively revise their intermediate features within the backbone for the prediction task. The overall architecture is shown in Fig. 6.

**Figure 6:**
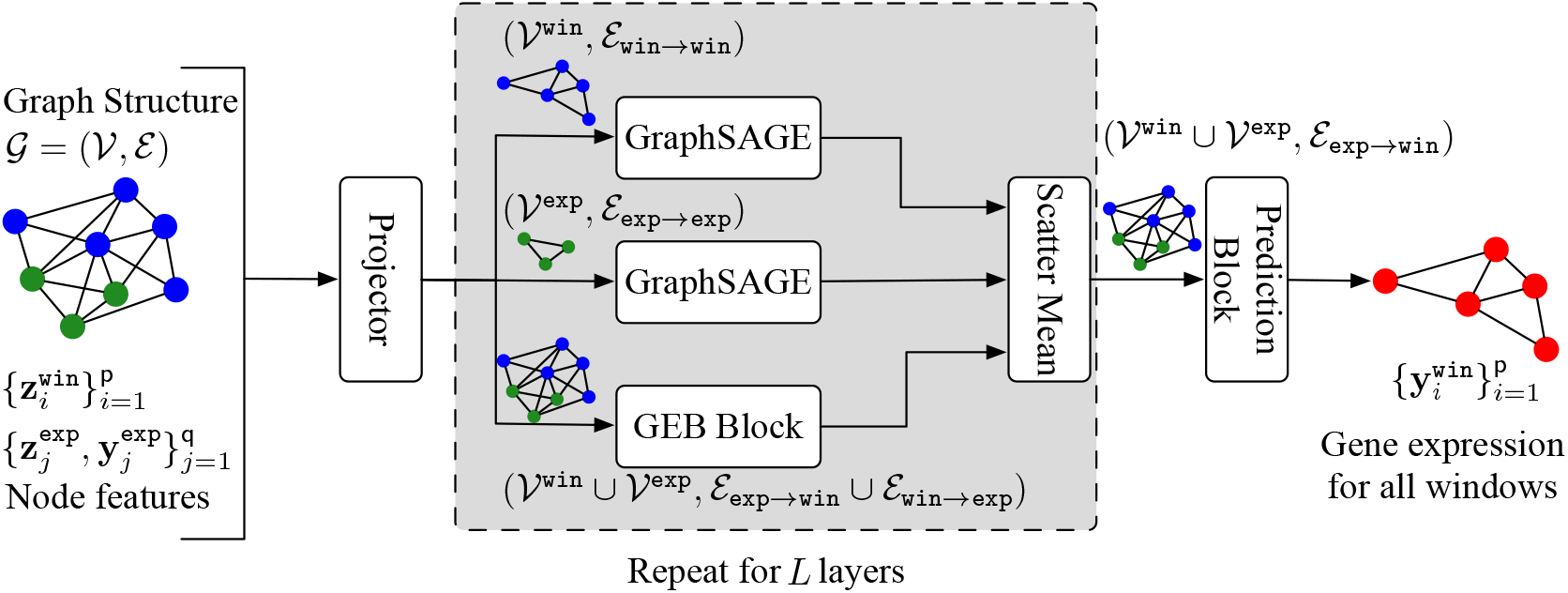
Architectures of our model. We have a graph 𝒢 = (𝒱, ℰ), where the node set 𝒱 = 𝒱^win^ ∪ 𝒱^exp^ is a union of windows and exemplars, and the edge set ℰ = ℰ_win→win_ ∪ ℰ_exp→exp_ ∪ ℰ_win→exp_ ∪ ℰ_exp→win_ contains four types of relations among nodes. We first refine features (i.e., 𝒢, 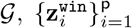 and 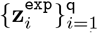 of each node by using a projector. We then respectively allow interactions among 𝒱^win^ and 𝒱^exp^ by two GraphSAGE block [9]. Meanwhile, we have a GEB block that revises the features of nodes contained in ^win^ and ^exp^ by considering edges between the two node sets. The scatter mean operation [41] average the features computed by the two GraphSAGE blocks and the GEB block for each node. Finally, after repeating the above operations for *L* layers, there is a prediction block that weights the contribution of each exemplar to a window for the gene expression prediction task. Finally, we have gene expression of all window nodes as 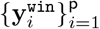. Note that the blue window nodes turn red, carrying gene expression prediction at each of the window nodes.

#### Projector

The features, 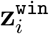 and 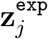, are embeded with a wide range of dataset-dependent attributes. We refine them to concentrate on the gene expression of interest by several multi-layer perceptrons (MLPs). Firstly, each window 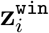 is projected by 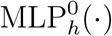. Secondly, for each 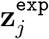, as its associated gene expression 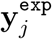, is available, we empower the refinement of 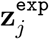 by 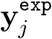. We concatenate 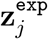 and 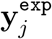 before feeding to 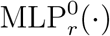. We have

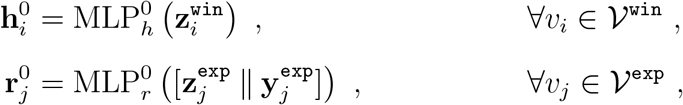

where the superscripts of 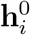 and 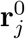 denote that they are initial refined feature, and · I · is a concatenation operator. 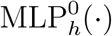 and 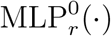 are three-layer perceptrons with LeakyReLU activation functions. The bottom two layers of 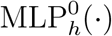 and 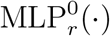 share the same parameter, restricting them in the same feature space.

#### Backbone

We use a sequence of GraphSAGE layers as our backbone [9], to allow interactions respectively among windows and exemplars. These interactions evolve each window feature under their neighborhood structural information and smooth each exemplar with neighbors to provide more accurate features that are used in the following layers. Assuming there is *L* layer and *l* ∈ [1, · · ·, *L*]. At *l*^th^ layer, we mathematically define

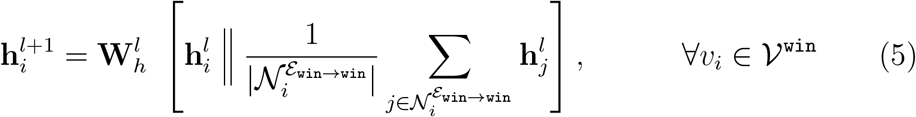

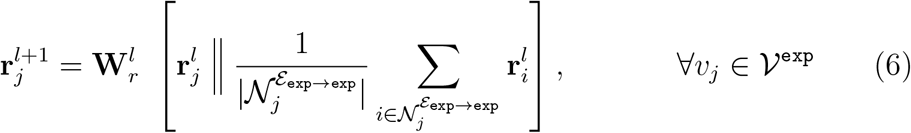

where 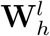 and 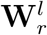 are linear weight matrices, and 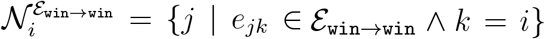 and 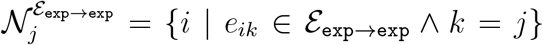 contain the index of neighborhood nodes under the given edge set.

#### GEB Block

This block is concurrent with the GraphSAGE backbone, bringing knowledge about gene expression 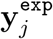 of the exemplar to the window feature 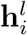. For brevity, we do not differentiate between the outputs of the GraphSAGE layer and the GEB block. At *l*^th^ layer, we project 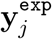 to 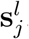, and have interactions between 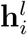 and 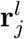 to obtain the revised window and exemplar features, i.e., 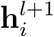 and 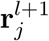.

In detail, the difference between 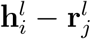 is passed to 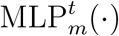, querying the potential knowledge of their gene expression difference. We chunk the outputs to **m**_*h,j,i*_ and **m**_*r,i,j*_. Then, they are used to retrieve gene expression from 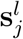 (i.e., a projection of 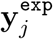), and adjust feature significance of 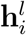 and 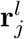. Semantically, **m**_*h,j,i*_ summarises ‘the existing knowledge of gene expression’, and **m**_*r,j,i*_ tells ‘the desired gene expression knowledge’. Mathematically, we have

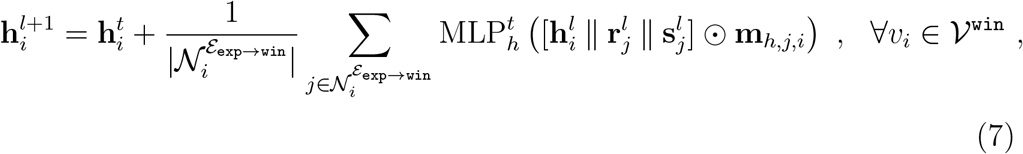

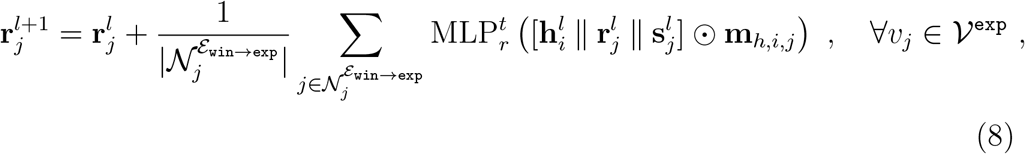

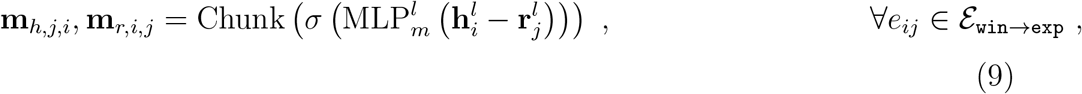

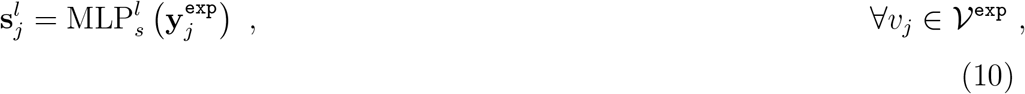

where 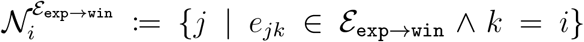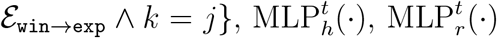, and 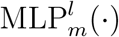 are multi-layer perceptrons, the ⊙ is element-wise multiplication, 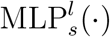 is a single layer perceptron, the Chunk(·) operator equally splits the input into two outputs, and *σ*(·) is a Sigmoid function. By scaling the magnitudes of the features in Eq. 7, we directly inject the knowledge about the gene expression of the exemplars into the window feature, while we also update the exemplar feature to facilitate both interactions with the exemplars and the window feature in following layers. We finally apply a scatter mean operation to aggregate the updated features from the GEB block and the GraphSAGE layers.

#### Prediction Block

We extend the attention pooling from [42] to consider the contribution of exemplars to a window toward the gene expression prediction, while also encouraging the model to refine exemplars feature in a more window-dependent way. We measure the importance **â**_*i,j*_ of each exemplar by using the difference between a window 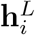 and a linearly projected exemplar 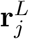. Then, the importance score of exemplars for a given window is normalized in the same vein with [43]. The normalized importance score **a**_*i,j*_ is used to weigh the exemplar and aggregate it into the window feature before making the gene expression prediction. Overall, our prediction block is defined as

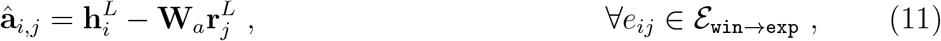

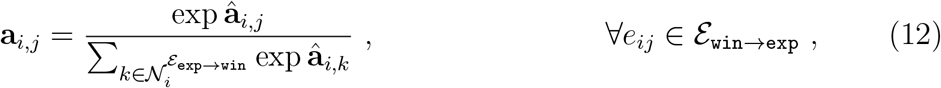

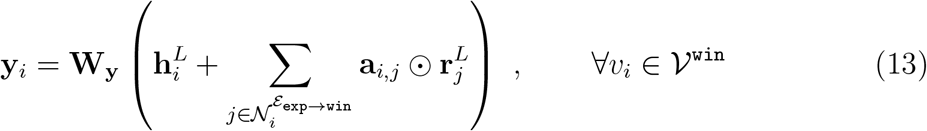

where **W**_*a*_ and **W**_**y**_ are linear weight matrix. Note that Eq. 12 performs computations element-wisely.

#### Objective

Our model is optimized with mean squared loss (i.e., the Euclidean distance) L_2_ and batch-wise PCC 𝓁_pcc_. We have

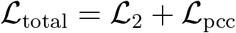

## 4. Experiments

### Datasets

We perform experiments in the publicly available STNet dataset [3] and 10xProteomic datasets^1^. The STNet dataset contains roughly 30,612 pairs of the slide image window and the gene expression. This dataset covers 68 slide images from 23 patients. Following [3], we target predicting expression of 250 gene types that have the largest mean across the dataset. The 10xProteomic dataset has 24,263 slide image windows and gene expression pairs from 6 slide images. We select target gene types in the same way as the STNet dataset. We apply log transformation and min-max normalization to the target gene expression. Our normalization method is different from [3] (they use log transformation and L_1_ normalization, i.e., log-transforming the division of the expression of each gene type by the sum of expression of all gene types, for each slide image window). Our normalization method allows independent analysis of expression prediction of each gene type.

### 4.1. Experimental Set-up

*Baseline Methods*. We compare with extensive SOTA methods in domains of gene expression prediction, ImageNet classification benchmarks, exemplar learning, and GCNs.

- STNet [3], NSL [4], and EGN [11]. They are the SOTA methods in gene expression prediction.
- ViT [43], MPViT [44] and CycleMLP [45]. We use the SOTA ImageNet classification methods in our task. They are strong baselines in our task. Specifically, we use ViT-B, MPViT-Base, and CycleMLP-B2. Please refer to [43, 44, 45] for details.
- Retro [22] and ViTExp. We explore the SOTA exemplar learning methods, where the input window is represented in image **w**_*i*_. However, Retro is originally developed for natural language processing. We adapt it by providing the feature extractor output as the exemplar features. ViTExp directly concatenates the exemplar features to the ViT patch representation. The exemplar features are added with an embedding to be differentiated from the patch representation of the slide image window. Both Retro and ViTExp are based on the ViT-B architecture [43].
- GraphSAGE [9], GATv2Conv [38], and TransformerConv [35]. In the same vein as the last categories, we adapt the SOTA GCNs for exemplar learning, by allowing communications from windows to windows, exemplars to exemplars, windows to exemplars, and exemplars to windows.

#### Evaluation Metrics

We evaluate the proposed methods and alternative baselines with PCC, mean squared error (MSE), and mean absolute error (MAE). We use PCC@F, PCC@S, and PCC@M to denote the first quantile, median, and mean of the PCC. The PCC@F verifies the least performed model predictions. The PCC@S and PCC@M measure the median and mean of correlations for each gene type, given predictions and ground truth for all of the slide image windows. Meanwhile, the MSE and MAE measure the sample-wise deviation between predictions and ground truth of each slide image window for each gene type. For PCC@F, PCC@S, and PCC@M, the higher value indicates better performance. In contrast, for MSE and MAE, the lower value means better performance.

#### Implementation Details

We use the ResNet18 [10] that is officially pretrained on the ImageNet-1K dataset as our feature extractor. We implement EGGN by using the *Pytorch* [46] and *PyTorch Geometric* [47] frameworks. EGGN is respectively trained from scratch for 50 epochs and 300 epochs on the STNet dataset and 10xProteomic dataset. We use batch size 1. All windows of a slide image composite a batch with size 1 in our case. We set the learning rate to 5 × 10^−4^. Our weight decay is 1 × 10^−4^. We use a GraphSAGE backbone with hidden dimensions 512 and layers 4. All the experiments are conducted with NVIDIA Tesla P100 GPUs.

#### Quantitative Evaluation

We compare our EGGN framework with the baselines on the STNet dataset and the 10xProteomic dataset (Tab. 1). As the gene expression prediction task emphasizes capturing the relativity variation, we bias on the PCC-related evaluation metrics, i.e., PCC@F, PCC@S, and PCC@M. Our EGGN consistently achieves the best performance in PCC@F, PCC@S, and PCC@M. Our findings are as follows: i) it’s worth noting that GCN-based approaches achieve higher PCC-related evaluation metrics than the approaches that predict gene expression individually from each window, *e*.*g*., EGN, the past SOTA method. GCN-based approaches are capable of modeling spatial relations among windows, capturing relative changes among window features which improve the PCC-related evaluation metrics; ii) GraphSAGE, GATv2Conv, and TransformerConv, the SOTA GCN methods, lead to the second-best performance in the STNet dataset and 10xProteomic dataset in PCC-related evaluations. Meanwhile, GraphSAGE finds the best MSE and MAE on the STNet dataset. These GCN models all outperform their counterparts, the SOTA methods in the ImageNet-1K classification task (*e*.*g*., CycleMLP and MPViT), evidencing our claims that reasoning spatial relations among windows out-weight feature learning in our gene expression prediction problem (Sec. 1); iii) the PCC@F of our model significantly outperforms the baseline methods. This metric evaluates the worst model capability by calculating the first quantile of PCC across all gene types. The majority of gene types covered by the first quantile have skewed expression distributions, which is the most challenging part of the prediction task. Our method has 0.012 - 0.024 higher than the second-best performance from other methods in PCC@F; iv) STNet and NSL fail to achieve good performance. Again, the gene expression-related features are usually non-uniformly distributed across the slide image. They predict the gene expression of each window independently, failing to capture feature dependency among the slide image. Moreover, NSL shows a negative correlation with PCC@F. This validates our claims that predicting gene expression directly from the color intensity is vulnerable, and it is only feasible in extreme cases, *e*.*g*., the example of tumor-related gene expression in Sec. 1.

**Table 1:** Quantitative gene expression prediction comparisons with SOTA methods on STNet dataset and 10xProteomic dataset. We bold the best results. We use ‘-’ to denote unavailable results. Models are evaluated by four-fold cross-validation and three-fold cross-validation on the above datasets. Our proposed EGGN framework consistently outperforms the SOTA methods in 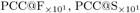 and 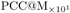 for both datasets. GraphSAGE finds the best 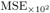 and 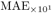 on the STNet dataset.

| Method | STNet Dataset |  |  |  |  | 10xProteomic Dataset |  |  |  |  |
| --- | --- | --- | --- | --- | --- | --- | --- | --- | --- | --- |
| | $MSE_{\times 10^2}$ | $MAE_{\times 10^1}$ | $PCC@F_{\times 10^1}$ | $PCC@S_{\times 10^1}$ | $PCC@M_{\times 10^1}$ | $MSE_{\times 10^2}$ | $MAE_{\times 10^1}$ | $PCC@F_{\times 10^1}$ | $PCC@S_{\times 10^1}$ | $PCC@M_{\times 10^1}$ |
| STNet [3] | 4.52 | 1.70 | 0.05 | 0.92 | 0.93 | 12.40 | 2.64 | 1.25 | 2.26 | 2.15 |
| NSL [4] | - | - | -0.71 | 0.25 | 0.11 | - | - | -3.73 | 1.84 | 0.25 |
| ViT [43] | 4.28 | 1.67 | 0.97 | 1.86 | 1.82 | 7.54 | 2.27 | 4.64 | 5.11 | 4.90 |
| CycleMLP [45] | 4.41 | 1.68 | 1.11 | 1.95 | 1.91 | 4.69 | 1.55 | 5.88 | 6.60 | 6.32 |
| MPViT [44] | 4.49 | 1.70 | 0.91 | 1.54 | 1.69 | 5.45 | 1.56 | 6.40 | 7.15 | 6.84 |
| Retro [22] | 4.53 | 1.71 | 0.99 | 1.74 | 1.79 | 5.25 | 1.65 | 5.46 | 6.35 | 6.04 |
| ViTExp | 4.46 | 1.69 | 0.87 | 1.72 | 1.74 | 5.04 | 1.66 | 5.59 | 6.36 | 6.00 |
| EGN [11] | 4.10 | 1.61 | 1.51 | 2.25 | 2.02 | 5.49 | 1.55 | 6.78 | 7.21 | 7.07 |
| GraphSAGE [9] | **3.79** | **1.57** | 1.98 | 2.90 | 2.83 | 4.98 | 1.65 | 6.82 | 7.42 | 7.25 |
| GATv2Conv [38] | 4.03 | 1.62 | 2.00 | 2.85 | 2.77 | 4.35 | 1.49 | 6.79 | 7.25 | 7.11 |
| TransformerConv [35] | 4.14 | 1.64 | 1.97 | 2.78 | 2.69 | 4.71 | 1.53 | 6.81 | 7.39 | 7.27 |
| Ours | 3.94 | 1.61 | **2.12** | **3.05** | **2.92** | **3.52** | **1.31** | **7.06** | **7.60** | **7.44** |

#### Quantitative Evaluation

We present the latent space visualization (Fig. 7), by considering the top performed models from Tab. 1. Fig. 7 (a, b, c) are models that predict gene expression of each window individually, and Fig. 7 (d, e, f) are best performed GCN-based methods. To enable a clean visualization, we randomly sample 256 slide image window features for each label, i.e., tumour and normal. Our method sufficiently separates the tumour features from the normal features.

**Figure 7:**
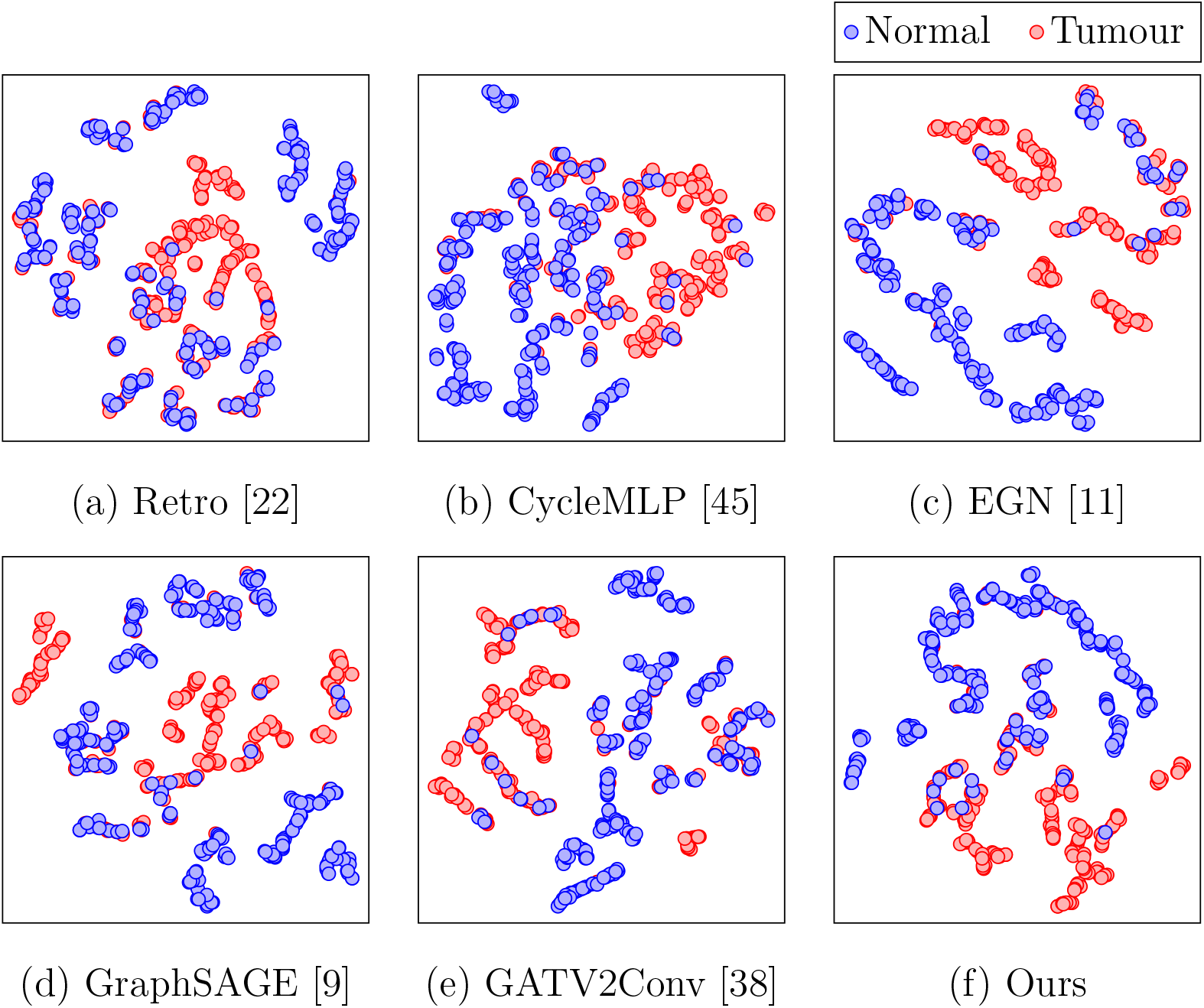
Quantitative evaluation of the top performed models from Tab. 1. We employ t-SNE [48] for feature dimension reduction. We use the extra labels (i.e., tumour and normal) from the STNet dataset for annotations.

### 4.2. Ablation study

We study the capability of each model component by conducting a detailed ablation study on the STNet dataset.

#### Number of Exemplars

We present PCC@M (Fig. 8 (a)), MSE (Fig. 8 (b)), and MAE (Fig. 8 (c)), by varying the number of exemplars used in our model from 1 to 15. Having 6 exemplars finds the best PCC@M, and we have the best MSE and MAE by using 3 exemplars. Again, our task emphasizes capturing relative gene expression changes. Thus, our final recommendation is to use 6 exemplars.

**Figure 8:**
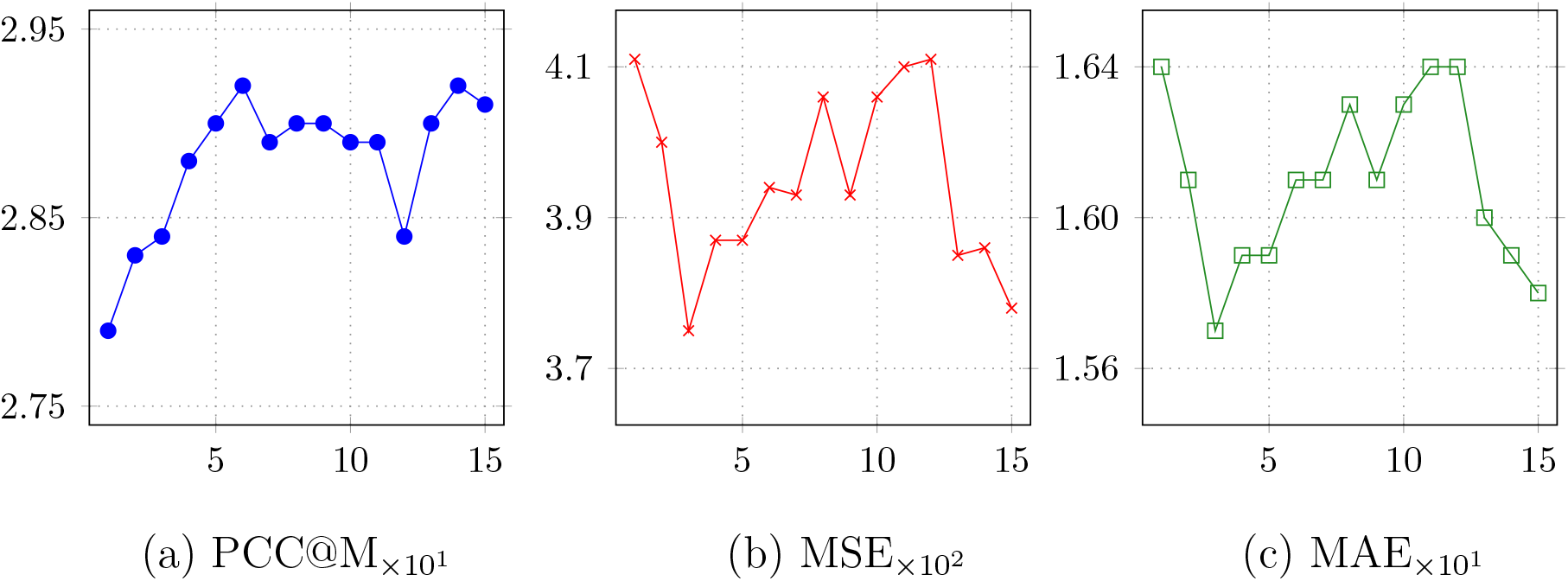
Ablation study on the number of exemplars used in our model. The number is varied from 1 to 15, and we respectively present PCC@M, MSE, and MAE in (a), (b), and (c).

#### Exemplar Retrieval

We explore alternative approaches for retrieving exemplars (Tab. 2). We study exemplar retrieval strategy by considering StyleEncoder [11], AlexNet [49], and ResNet18 [10]. Among the three backbone networks, StyleEncoder is learned in an unsupervised manner with slide images, while AlexNet and ResNet18 are trained with supervision for ImageNet-1K (i.e., natural RGB images) classification task. We explore diverse distance matrices including LPIPS [50], L_2_, L_1_, and *cosine* (i.e., cosine similarity). We have the following findings: i) LPIPS distance retrieved exemplars lead to bad model performance. The distance is trained based on the human perception of natural RGB images. These natural RGB images are different from the slide images, leading to bad exemplar retrieves, and damaging the model performance; ii) the StyleEncoder is optimized for reconstructing slide images. However, as shown in Fig. 3, gene expression-related knowledge is not balanced across the dataset, imposing biased knowledge to the StyleEncoder. When performing interactions among windows and exemplars in our model, such biases are potentially amplified and then decrease the model performance; iii) using ResNet18 with ℒ_1_ distance achieves the best performance, though the pre-trained ResNet18 lacks gene expression-related knowledge. This retrieval method is an expert at encoding knowledge of image textures, benefiting from the large-scale training dataset, ImageNet-1K for our task. This allows modeling the relation among windows and exemplars in a more accurate way.

**Table 2:** Ablation study on exemplar retrieval. We compare features from StyleEncoder, AlexNet and ResNet18 for exemplar retrieval. We use the ResNet18 with ℒ_1_ for exemplar retrieval in our EGGN framework.

| Feature Space | Distance Metrics | MSE$\times 10^2$ | MAE$\times 10^1$ | PCC@M$\times 10^1$ |
| --- | --- | --- | --- | --- |
| StyleEncoder | *cosine* | 3.87 | 1.59 | 2.61 |
| StyleEncoder | $\mathcal{L}_2$ | 4.05 | 1.62 | 2.60 |
| StyleEncoder | $\mathcal{L}_1$ | 4.02 | 1.62 | 2.62 |
| AlexNet | LPIPS | 4.21 | 1.65 | 2.64 |
| ResNet18 | *cosine* | 4.00 | 1.62 | 2.88 |
| ResNet18 | $\mathcal{L}_2$ | 4.03 | 1.62 | 2.86 |
| ResNet18 | $\mathcal{L}_1$ | 3.94 | 1.61 | 2.92 |

#### Model Architectures

We study the performance of three baseline settings, ‘Backbone only’, ‘w/o EB block’, and ‘w/o projector’, in Tab. 3. The ‘Backbone only’ setting uses the GraphSAGE backbone architecture, considering the windows from the graph only. The ‘w/o GEB block’ setting removes the GEB block, and replaces our prediction block with a single linear layer for gene expression prediction. The ‘w/o projector’ setting replaces the projector with a linear layer to unify the dimension. Our findings are as follows: i) the ‘Backbone only’ setting achieves the worst performance because of the absence of extra knowledge from the exemplar; ii) the ‘W/o GEB’ block has the second-worst performance because of a similar reason. However, it has a projector to refine the ResNet18 feature to gene expression-related features; iii) with all proposed components, we have the best performance.

**Table 3:** Ablation study on model architectures.

| Settings | $\text{MSE}_{\times 10^2}$ | $\text{MAE}_{\times 10^1}$ | $\text{PCC@M}_{\times 10^1}$ |
| --- | --- | --- | --- |
| Backbone only | 4.22 | 1.66 | 2.76 |
| W/o GEB | 4.18 | 1.65 | 2.85 |
| W/o Projector | 4.10 | 1.63 | 2.84 |
| Ours | 3.94 | 1.61 | 2.92 |

#### Graph Construction

We study if constructing the exemplar nodes in our graph can generally benefit other baseline GCN frameworks, i.e., Graph-SAGE, GATv2Conv, and TransformerConv (Tab. 4). As shown, the performance of these models has consistently been improved, by leveraging extra knowledge from the exemplars. With our GEB blocks, we have the best performance.

**Table 4:** Ablation study on graph construction. The performance of our method in the ‘w/o exemplars’ setting is omitted, as our method is designed for exemplar learning only.

| Settings | Method | $\text{MSE}_{\times 10^2}$ | $\text{MAE}_{\times 10^1}$ | $\text{PCC@M}_{\times 10^1}$ |
| --- | --- | --- | --- | --- |
| w/o<br>exemplars | GraphSAGE | 4.22 | 1.66 | 2.76 |
|  | GATv2Conv | 4.34 | 1.68 | 2.75 |
|  | TransformerConv | 4.45 | 1.70 | 2.62 |
| w/<br>exemplars | GraphSAGE | 3.79 | 1.57 | 2.83 |
|  | GATv2Conv | 4.03 | 1.62 | 2.77 |
|  | TransformerConv | 4.14 | 1.64 | 2.69 |
|  | Ours | 3.94 | 1.61 | 2.92 |

## 5. Limitation and Future Work

### Limitation

Our method is capable to predict the gene expression of multiple windows in a single forward pass. However, computation costs have to be wasted, when a user is only interested in the gene expression of one window in a slide image. With a single window, our model is equivalent to sequential stacking linear layers, lacking interactions among windows and exemplars, and decreasing the model performance. Thus, dummy windows from the slide image are needed to be sampled for forming a graph. The sampling and prediction processes consume extra computation resources.

### Future Work

Though exemplars could be used as anchors for allowing information propagation for windows distributed in a slide image, a more explicit solution for facilitating information propagation needs to be explored. In our future work, we will study downsampling and upsampling operations for our exemplar and window-based graphs to build an encoder and decoder-based architecture to allow broader information propagation in the slide image.

## 6. Conclusion and Broader Impact

This paper proposes an EGGN framework to accurately predict gene expression from each fine-grained area of tissue slide image, i.e., different windows. EGGN uses the GraphSAGE as a backbone while integrating with exemplar learning concepts. We first have an extractor to retrieve the exemplars of the given tissue slide image window. Then, we construct a graph to connect windows within the same slide image and their corresponding exemplars, and propose a GEB block to progressively revise the intermediate GraphSAGE feature by reciprocating with the nearest exemplars. With extensive experiments, we demonstrate the superiority of the EGGN framework over the SOTA methods. EGGN is promising to facilitate studies on diseases and novel treatments with accurate gene expression prediction.

## Acknowledgement

The authors would like to thank Machine Learning & Artificial Intelligence Future Science Platforms, CSIRO for computation resource funding.

## Footnotes

1 https://www.10xgenomics.com/resources/datasets

